# A non-ligand surrogate agonist antibody that enhances canonical Wnt signaling and bone regeneration

**DOI:** 10.1101/2022.09.25.509440

**Authors:** Nam-Kyung Lee, Scott Bidlingmaier, Yang Su, Youngho Seo, Bin Liu

**Affiliations:** Department of Anesthesia, UCSF Helen Diller Family Comprehensive Cancer Center, University of California at San Francisco, 1001 Potrero Ave., 1305, San Francisco, CA 94143-1305; Department of Radiology and Biomedical Imaging, University of California at San Francisco, 185 Berry Street, San Francisco, CA 94107

**Keywords:** Wnt/β-catenin signaling, agonist antibody, non-ligand surrogate, human monoclonal antibody, LRP6, R-spondin, bone formation, cancer-induced bone loss, multiple myeloma

## Abstract

The Wnt signaling pathway promotes tissue regeneration and is a promising therapeutic target for treatment of osteolytic bone diseases. Here we report the discovery of a novel type of canonical Wnt agonist antibody that does not operate as a ligand surrogate. The antibody increases Wnt/β-catenin signaling with or without exogenously provided Wnt ligands. It binds to a site on the P3 domain of LRP6 that is distinct from where the Wnt3a ligand and the DKK1 antagonist bind. The agonist effect persists in the presence of DKK1 and is further amplified by R-spondin even when Wnt ligands are not provided, suggesting a potential use for this antibody in ligand-low or insufficient settings. The antibody induces osteoblastic differentiation and mineralization *in vitro* and restores bone loss *in vivo* in a myeloma-derived intrafemoral mouse model, opening a potential path for therapeutic development in osteolytic diseases caused by cancer and aging.

## Introduction

The canonical Wnt/β-catenin signaling pathway is involved in various biological processes including tissue regeneration, stem cell regulation, cell proliferation and differentiation (Clevers et al., 2014; Lien and Fuchs, 2014; Steinhart and Angers, 2018). The critical role of canonical Wnt signaling in driving bone formation has been shown by several studies (Baron and Kneissel, 2013; Florio et al., 2016; Liu et al., 2016; McDonald et al., 2017; Pozzi et al., 2013). The importance of canonical Wnt signaling in bone formation is also shown by bone degenerative effects of Wnt signaling inhibitors such as sclerostin (SOST) or Dickkopf Wnt signaling pathway inhibitor 1 (DKK1). An anti-SOST monoclonal antibody (romosozumab) has been approved for osteoporosis treatment in postmenopausal women (Markham, 2019). In addition, the anti-DKK1 antibody BHQ880 has been clinically evaluated for restoration of osteolytic bone loss driven by multiple myeloma (Fulciniti et al., 2009; Iyer et al., 2014; Munshi and Anderson, 2013).

Although the aforementioned approaches targeting inhibitory ligands are achieving promising results in the clinic, there is still a need to develop additional effective therapies. Therapies targeting inhibitors may be less effective if Wnt ligands are absent or below a critical threshold in the disease region. In addition, the anti-inhibitor approach is limited to the particular inhibitor that a monoclonal antibody is designed to bind and neutralize. For example, while romosozumab blocks SOST, it does not block DKK1, resulting in potentially limited blocking of inhibitory activities toward Wnt signaling (Joiner et al., 2013).

An alternative approach is to directly activate Wnt signaling using a canonical Wnt pathway agonist. Canonical Wnt signaling is induced by two distinct Wnt co-receptors, the G protein-coupled receptor Frizzled (Fzd) and low-density lipoprotein receptor-related protein 5 or 6 (LRP5/6). Binding of Wnt ligands drives the formation of the Fzd-Wnt-LRP6 complex that leads to LRP6 phosphorylation to initiate the signaling. Inhibition of canonical Wnt signaling by anti-LRP6 antibodies has been reported (Ettenberg et al., 2010). A ligand surrogate-based Wnt agonist capable of activating Wnt signaling and promoting bone formation has been reported (Janda et al., 2017) which consists of an anti-Fzd scFv and the DKK1 LRP6-binding domain, thereby mimicking the mechanism of natural Wnt ligands. Another ligand surrogate-based Wnt agonist has also been reported that consists of an anti-FZD scFv and an anti-LRP6 single domain antibody (Fowler et al., 2021). Other Wnt ligand surrogates have also been described (Chen et al., 2014; Tao et al., 2019) that explore multivalency and crosslinking to enhance signaling, a mechanism that may also be used by natural Wnt ligands and co-activators. Fundamentally, however, those ligand surrogates compete with endogenous Wnt ligands for binding to receptor complex and are subject to inhibition by endogenous inhibitors such as DKK1 and SOST that bind to ligand binding sites.

We hereby describe the discovery of a different type of canonical Wnt pathway agonist that does not follow the ligand surrogate mode. Using an unbiased functional screening where we selected phage antibody display libraries on LPR6, we identified a novel human monoclonal antibody with agonist activity. We showed that this agonist antibody binds to a distinct site on LRP6 that does not overlap with where Wnt ligands and inhibitors bind, and acts as a novel canonical Wnt pathway agonist that does not operate as a Wnt ligand surrogate. It activates canonical Wnt signaling even when no Wnt ligands are provided, and the activation is further amplified by R-spondin. Furthermore, the agonist activity is not blocked by endogenous inhibitors that bind to LRP6. This novel agonist antibody is effective in promoting osteoblastic differentiation *in vitro* and restoring *in vivo* bone loss caused by intrafemorally implanted multiple myeloma cells in mice. Thus we have demonstrated that there is a class of novel canonical Wnt pathway agonist antibodies that bind to LRP6 at sites distinct from where Wnt ligands or inhibitors bind and do not follow the ligand surrogate mode of action. The demonstrated *in vivo* bone restoring activity suggests that this type of agonist antibodies have potential to be developed into therapeutics for treatment of pathological bone loss and perhaps other degenerative conditions caused by insufficient canonical Wnt signaling.

## Materials and Methods

### Cell lines

Human embryonic kidney (HEK) 293A and 293 cell lines, multiple myeloma MM1.S cell line, mouse L Wnt-3a cell line, mouse pre-osteoblast MC3T3-E1 cell line, mouse mesenchymal C3H/10T1/2 cell line were obtained from American Type Culture Collection (ATCC). Cells were maintained in DMEM, RPMI1640, or α-MEM supplemented, per vendor instructions, with 10% fetal bovine serum (FBS, Fisher Scientific) and 100 μg/ml penicillin/streptomycin (Axenia BioLogix) at 37°C in a humidified atmosphere containing 5% CO_2_. Myeloma cell line-derived conditioned medium (CM) was collected by centrifugation of supernatant from cell culture at 70~80% confluency. All cell lines were used within six passages and were not authenticated by short tandem repeat profiling. Cells were tested negative for Mycoplasma using PCR Mycoplasma detection kit (abm, Canada).

### Selection of Wnt agonist antibodies from phage display libraries

Recombinant LPR6 P3E3P4E4 domain was produced as a Fc fusion protein and purified on protein A column as previously described (Lee et al., 2018). Naïve phage antibody display libraries were selected on biotin-labeled LRP6-P3E3P4E4 fragment as previously described (Lee et al., 2018). After three rounds of selection, monoclonal phage were arrayed into 96-well plates and tested for binding to LRP6-transfected HEK293 cells by flow cytometry. Unique scFv antibodies were identified by sequencing from LRP6-binding phages, and individual phage clones were amplified and purified for further characterization.

### Plasmids, cloning, and site-directed mutagenesis

Full-length human LRP6 was cloned into pCMV-Entry (Origene) and used for sub-cloning, point mutation, or transfection. Truncation constructs of LRP6 ectodomains were generated by cloning into pCMV-Entry and used for transient expression. Alanine mutants of LRP6 ectodomain were generated using QuikChange Site-Directed Mutagenesis Kit (Agilent Technologies) according to the manufacturer’s protocol. The vector pFUSE-hIgG1-Fc2 (InvivoGen) was used for cloning Fc-fusion constructs. To construct IgG, variable heavy (VH) and kappa light (Vκ) chain genes were sub-cloned into Abvec Ig-γ and Ig-κ plasmids kindly provided by Dr. Patrick Wilson at University of Chicago (Smith et al., 2009) with modifications (Lee et al., 2018). For Fab constructs, CH2-CH3 was deleted from Ig-γ Abvec and a hexahistidine tag was introduced at the C-termini of CH1. The T-cell factor/lymphoid enhancer factor (TCF/LEF) luciferase reporter SuperTopFlash (STF) and the control pRL-SV40 Renilla luciferase constructs (Addgene) were used for Wnt/β-catenin-responsive reporter assays. Wnt ligands were provided by transient transfection of the pcDNA-Wnt1 or - Wnt3a expression plasmid (Addgene) or as recombinant products (R&D Systems).

### Production of recombinant proteins and antibodies

For transient transfection, plasmid DNA was resuspended in Opti-MEM (Life Technologies), mixed with polyethylenimine, and added to HEK293A cells. 24 h post transfection, media was changed to Freestyle 293 expression medium (Gibco) and the cells were further cultured for 6-8 days. Secreted proteins in supernatants were collected, filtered, and purified on protein A agarose (Thermo Scientific) for Fc-fusions and IgGs or Ni-NTA resin (Thermo Scientific) for Fab according to the manufacturer’s protocols.

### SuperTopFlash (STF) Luciferase reporter assays

Cells cultured in a 24- or 96-well plate were transiently transfected with STF luciferase reporter and the internal control pRL-SV40 Renilla luciferase expression plasmid using TransIT-2020 (Mirus Bio), with or without Wnt1- or Wnt3a-expression construct as indicated in Results. To express LRP6 truncates and alanine mutants, plasmid DNA encoding the constructs was co-transfected with reporter plasmids into HEK293 cells. The 6-6 antibody diluted in culture medium was added to the transfected cells, with or without recombinant DKK1 or RSPO2 (R&D Systems) as indicated in Results, and further incubated for 16 h. Firefly luciferase and Renila luciferase activities were detected using Dual-Luciferase Reporter Assay System (Promega) and normalized as described previously (Lee et al., 2018). Data were expressed as a fold relative to a control group transfected only with reporter constructs.

### Apparent K_D_ determination

The apparent K_D_ of antibodies was analyzed by flow cytometry as described (Lee et al., 2018). Briefly, cells were trypsinized, washed, and resuspended in PBS with 1% FBS. Antibodies serially diluted in PBS/1% FBS were incubated with target cells (3 x 10^5^ cells/tube) overnight at 4 °C. Cells were washed, incubated with Alexa Fluor^®^647-labeled goat anti-human IgG (Jackson ImmunoResearch) for 1 h at 4 °C, washed three times with PBS and analyzed using a BD Accuri C6 flow cytometer (BD Biosciences). Median Fluorescence Intensity (MFI) values were analyzed by curve fitting (GraphPad) to determine the apparent affinity.

### Bio-layer Interferometry

Competitive binding activity between anti-LRP6 Fab (6-6 Fab) and recombinant Wnt3a or DKK1 to LRP6 ectodomain was studied by bio-layer interferometry (BLI) using a BLItz (ForteBio) instrument. Protein A biosensors (ForteBio) were loaded with human LRP6-ECD-Fc (R&D Systems) for 120 sec, and dipped in recombinant Wnt3a for 75 sec followed by a mixture of Wnt3a + 6-6 Fab, or Wnt3a + DKK1 for 75 sec. Baselines were determined for 30 sec before and after the loading step according to the manufacturer’s instructions.

### Antibody-receptor docking analysis

Antibody variable fragment (Fv) consisting of 6-6 VH and Vκ sequences was generated by homology modeling using RossettaAntibody (Weitzner et al., 2017). Docking models between the Fv and LRP6-P3E3 domain obtained from 3S8Z (Cheng et al., 2011) or 4A0P (Chen et al., 2011) were generated using ZDOCK (Pierce et al., 2014). The DKK1-LRP6-P3 interaction is based on published structural study (Cheng et al., 2011). Wnt3a-, DKK1- or 6-6 Fv-binding residues in docking models were analyzed and visualized using the PyMOL Molecular Graphics System (Schrödinger, LLC).

### Osteogenic differentiation

Differentiation of C3H10T1/2 cells was induced as described previously (Zhong et al., 2016). In brief, C3H10T1/2 cells were cultured in normal growth medium (α-MEM, 10% FBS, 100 μg/ml penicillin/streptomycin) in 24-well culture plates at 70~80 % confluency. The following day, the culture medium was changed to osteogenic medium (growth medium supplemented with 50 μg/ml ascorbic acid, 10 mM β-glycerol phosphate) and replaced with osteogenic medium every 2-3 days. Cells were either cultured in the osteogenic medium, or treated with 30% L cell Wnt3a-conditioned medium (Wnt3aCM) or Wnt3aCM plus 6-6 for 21 days. To determine matrix mineralization, cells were stained using Alizarin Red S (ARS) Staining Quantification Assay (ScienCell Research Laboratories), and images were taken using BIOREVO BZ-9000 microscope (Keyence). ARS dyes were extracted from the stained cells and quantified according to manufacturer’s instructions.

### Quantitative real-time PCR (qRT-PCR)

Osteogenic differentiation of C3H/10T1/2 cells was induced as described above for 3 days. Total RNA was isolated using TRIzol™ reagent (Invitrogen) and used to generate cDNA using High-Capacity cDNA Reverse Transcription Kit (Applied Biosystems) according to the manufacturer’s protocol. qRT-PCR was performed with 15 ng of cDNA using Power SYBR Green PCR Master Mix (Applied Biosystems) on the ABI 7300 real time PCR system (Applied Biosystems). All reactions were conducted in duplicate and copy numbers for a target gene transcript were normalized to Glyceraldehyde 3-phosphate dehydrogenase (GAPDH). Data are presented as the relative mRNA expression in antibody-treated vs. untreated control cells. Specific primer sets for each target gene were as follows; GAPDH-F: 5’-GGCCTCACCCCATTTGATGT-‘3, GAPDH-R: 5’-CATGTTCCAGTATGACTCCACTC-‘3, ALP-F: 5’-AACCCAGACACAAGCATTCC-‘3, ALP-R: 5’-GCCTTTGAGGTTTITGGTCA-‘3, RUNX2-F: 5’-GAATGGCAGCACGCTATIAAATCC-‘3, RUNX2-R: 5’-GCCGCTAGAATICAAAACAGTIGG-‘3, BMP2-F: 5’-GGGACCCGCTGTCTTCTAGT-‘3, BMP2-R: 5’-TCAACTCAAATTCGCTGAGGAC-‘3, OC-F: 5’-CTGACCTCACAGATGCCAA-‘3, OC-R: 5’-GGTCTGATAGTCTGTCACAA-‘3.

### Alkaline phosphatase (ALP) activity assay

Cells were incubated with the 6-6 agonist antibody and Wnt3aCM (30% by final volume) for 7 days in the osteogenic medium as described above, washed, harvested, and lysed by repetitive freezing-thawing cycles in NP-40 buffer (150 mM NaCl, 1.0% NP-40, 50 mM Tris, pH 8.0) supplemented with protease inhibitors (Cell Signaling Technology). ALP activity in cell lysates was measured using *p*-nitrophenyl phosphate (Sigma-Aldrich) according to manufacturer’s instructions. ALP activity was normalized against the control group without 6-6 and Wnt3a treatment.

### *In vivo* micro-CT scanning

The micro x-ray computed tomography (micro-CT), a component of VECTor4/CT (MILabs B.V., Utrecht, The Netherlands) preclinical imaging system was used for *in vivo* bone scan. In order to visualize the femur and its joints, the field of view of micro-CT was set around the femur using built-in optical cameras, followed by CT acquisition with x-ray tube settings of 50 kVp and 0.24 mA. A total of 1,440 projections over 360° were acquired in a step-and-shoot mode with x-ray exposure time of 75 ms at each step. No data binning was applied during acquisition (i.e., 1×1 binning). During the CT data acquisition, animals were kept under anesthesia using isoflurane (approximately 2% isoflurane mixed with medical-grade oxygen). Image reconstruction after the projections were acquired was performed using the vendor-provided conebeam filtered backprojection algorithm. The reconstructed image volumes were in the voxel size of 0.02 mm × 0.02 mm ×0.02 mm. The volumetric matrix sizes were dependent on the field of view selected during the reconstruction step only focusing on distal femurs. After the reconstruction, the image volumes were processed to show the common orientation by re-orienting the isotropoic volumes using PMOD (PMOD Technologies, Zurich, Switzerland).

### *In vivo* bone restoration study

2 x 10^5^ MM1.S cells were implanted in the right femur of NOD/SCID/IL-2Rγ^-/-^(NSG) female mice. One week later, mice were randomized (n = 5/group) and treated intraperitoneally with the vehicle (PBS) or 6-6 IgG at 10 mg/kg every week for a total of 6 injections. One week post treatment, mice were anesthetized and scanned by micro-CT. One week post CT scanning, blood and femurs were collected from the mice, and free human Ig-lambda light chains in serum was assessed using a Human Lambda ELISA Kit (Bethyl Laboratories) according to manufacturer’s instructions. All mouse studies were performed according to UCSF Institutional Animal Care and Use Committee-approved protocols.

### Trabecular and cortical bone image analysis

CT data files were used for 3D reconstruction and generating planar images of whole, distal, and proximal regions of the femurs using BoneJ2 plug-in operated by Fiji software as described (Doube et al., 2010; Schindelin et al., 2012). The micro-architectural parameters of trabecular and cortical bones were analyzed using stacked 3D bone images, and bone volume over tissue volume (BV/TV), trabecular bone thickness (Tb.Th), and cortical thickness (Ct.Th) were obtained.

### Histology and immunohistochemistry

Femurs were dissected to remove soft tissue, fixed in 10% neutral-buffered formalin, and decalcified in 14% EDTA for 4 weeks. The tissues were embedded in paraffin and cut into 4 μm sections, stained with hematoxylin and eosin (H&E) or anti-human Ig-lambda light chain antibody (Abcam) using methods described (Su et al., 2018). Stained sections were imaged using a BIOREVO BZ-9000 microscope (Keyence).

### Statistical analysis

For two-group comparisons, two-tailed Student’s t-test was used, and P < 0.05 was used to reject the null hypothesis. For multiple (three or more) group comparisons, the one-way analysis of variance (ANOVA) was used using the Tukey’s test.

## Results

### Identification of a novel human monoclonal antibody that binds the P3 domain of LRP6 and activates canonical Wnt signaling

To identify novel LRP6-binding Wnt pathway agonist antibodies, we first generated a recombinant fragment of the extracellular domain of LRP6, specifically the P3E3P4E4 domain. We next selected phage human antibody display libraries against this LRP6 fragment, and identified binding clones. These LRP6-binding clones were tested for agonist effect on canonical Wnt signaling using the SuperTopFlash (STF) reporter assay on HEK293 cells transfected with Wnt ligand expression plasmids. One antibody, 6-6, was identified to have agonist activity as a phage and was converted into a human IgG. The apparent affinity of 6-6 IgG for LRP6 was measured on HEK293 cells and found to be ~ 5 nM (Supplemental **Figure S1A**). The agonist activity was re-tested using the 6-6 IgG on HEK293 cells expressing Wnt ligands (Wnt3a or Wnt1). As shown in **Figure 1A**, 6-6 IgG showed agonist effects for both Wnt3a and Wnt1 mediated signaling. Interestingly, the 6-6 IgG showed agonist activity even in the absence of exogenously added Wnt ligands (**Figure 1B**). We performed a titration experiment using 6-6 and recombinant human Wnt3a (rhWnt3a) and found that both 6-6 (**Figure 1C**) and rhWnt3a (**Figure 1D**) activate canonical Wnt signaling in a concentration dependent manner (half-maximal effective concentration (EC50) values are about 2-4 nM for 6-6 vs. 18 nM for rhWnt3a). Furthermore, the agonist effect of 6-6 is additive to that of rhWnt3a (**Figure 1E**).

**Figure 1.**
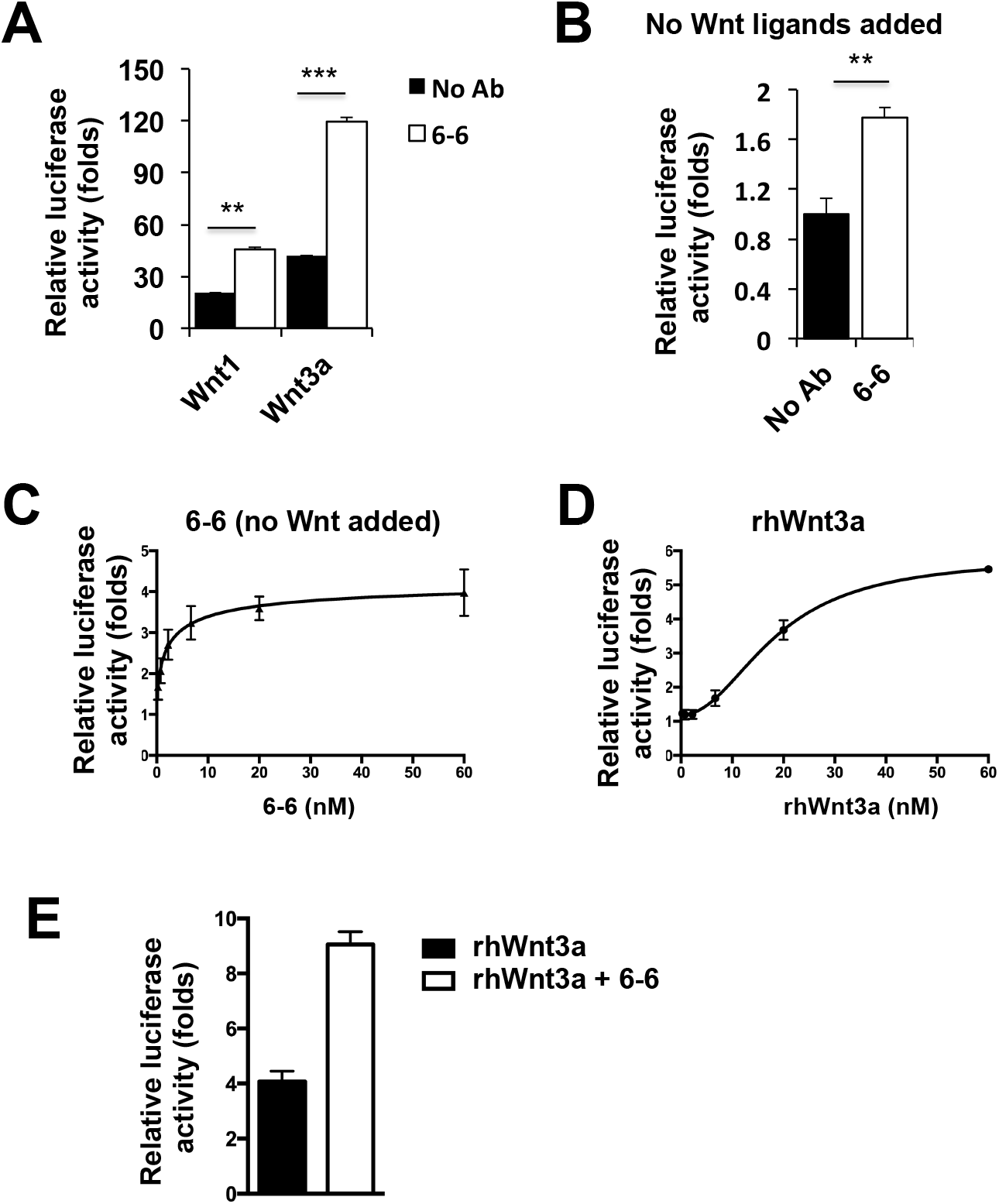
Novel Wnt-agonist human monoclonal antibody binding to the P3 domain of LRP6. (**A**) Activation of canonical Wnt signaling by 6-6. HEK293 cells transfected with the STF reporter and Wnt ligand expression constructs were incubated with or without 6-6 IgG (100 nM). Error bars represent SD for n = 2. ** P < 0.01; *** P < 0.001. (**B**) 6-6 agonist activity when no Wnt ligands are provided. HEK293 cells transfected with the STF reporter were incubated with or without 6-6 IgG (100 nM), and luciferase activity was normalized against a control without antibody treatment. Error bars represent SD for n = 2. ** P < 0.01. (**C**) STF reporter assay using HEK293 cells transfected with the STF reporter over a range of 6-6 IgG concentrations. Error bars represent SD for n = 3. EC50 is estimated by curve fitting to be 2.12 ± 2.56 nM. (**D**) STF reporter assay using HEK293 cells transfected with the STF reporter over a range of rhWnt3a concentrations. Error bars represent SD for n = 3. EC50 is estimated by curve fitting to be 18.62 ± 1.19 nM. (**E**) Additive effect of 6-6. The STF reporter assay was performed on HEK293 cells transfected with the STF reporter and exposed to rhWnt3a (20 nM), or rhWnt3a (20 nM) plus 6-6 IgG (20 nM). Error bars represent SD for n = 3. The relative luciferase activity (fold over untreated HEK293 cells transfected with the STF reporter) is 4.08 ± 0.38 for rhWnt3a and 8.93 ± 1.48 for rhWnt3a plus 6-6 IgG.

This novel canonical Wnt pathway agonist 6-6 antibody does not compete with a previously identified LRP6 P3E3 binder (Supplemental **Figure S1B**) that is antagonistic to Wnt/β-catenin signaling and competes with ligand binding (Lee et al., 2018). To map where 6-6 binds to LRP6, we first generated a series of truncation mutants of the LRP6 P3E3P4E4 domain (**Figure 2A**), expressed them in HEK293 cells by transient transfection and studied 6-6 binding by flow cytometry. We found that 6-6 binds to the P3 domain of LRP6 as deletion of P3 but not other segments caused a complete loss of 6-6 binding (**Figure 2B**). We further studied 6-6 agonist activity on LRP6 truncation mutants using the STF reporter assay (**Figure 2C**). The 6-6 IgG activated canonical Wnt signaling in cells expressing the LRP6 full-length and LRP6 P3E3P4E4 constructs but not other variants where the P3 domain is deleted, consistent with results from the cell binding study showing that 6-6 binds to the P3 domain.

**Figure 2.**
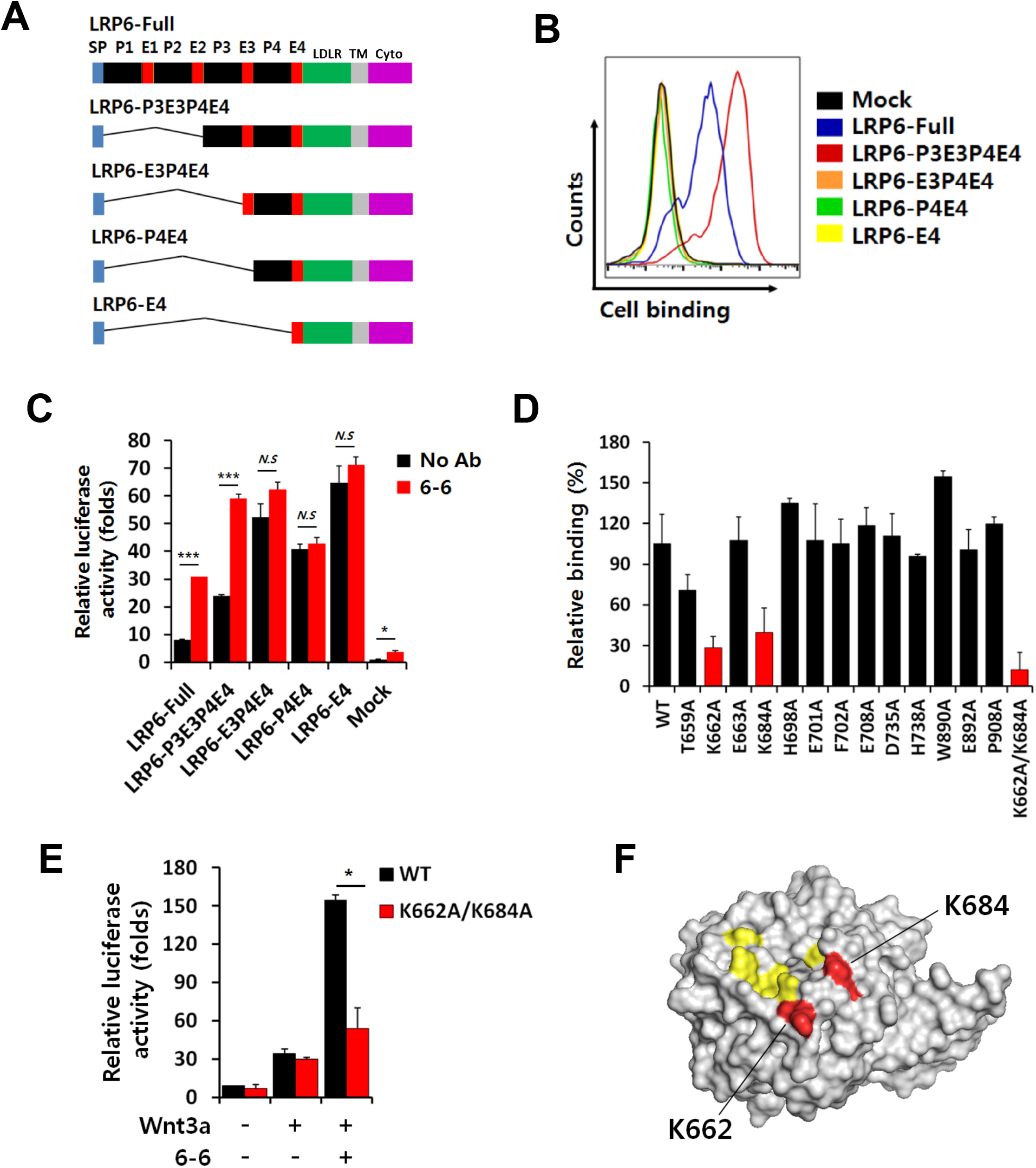
6-6 binds to a unique site on LRP6 that does not overlap with where Wnt ligand binds. (**A**) Deletion mapping of 6-6 binding sites on LRP6. Deletion constructs studied are shown. SP: Signal peptide; P1, 2, 3 and 4: Beta-propeller domains 1, 2, 3 and 4, respectively; E1, 2, 3 and 4: EGF-like domains 1, 2, 3 and 4, respectively; LDLR: Low-density lipoprotein receptor type A domain; TM: Transmembrane domain; Cyto: cytoplasmic domain. (**B**) Analysis of 6-6 binding to truncated cell surface LRP6. Each LRP6 truncation plasmid was separately transfected into HEK293 cells with a green fluorescent protein (GFP)-expressing plasmid. Binding of 6-6 IgG in GFP-positive cell population was analyzed by flow cytometry. (**C**) Assessment of Wnt-agonist activity of 6-6 IgG on cells expressing truncated LRP6. HEK293 cells were transfected with LRP6 truncation constructs and the STF reporter plasmid and incubated in Wnt3aCM with or without 6-6 IgG. Error bars represent SD for n = 2. *** P < 0.001. n.s.: not significant. (**D**) Fine epitope mapping by alanine scan. LRP6 single mutants and the double mutant (K662A/K684A) were separately transfected into HEK293 cells. An anti-LRP6 scFv-Fc fusion that binds to the LRP6-P1 domain was used as a control to confirm cell surface expression of LRP6. Binding of 6-6 was determined by flow cytometry, and median fluorescence intensity (MFI) values were normalized against MFI of the P1-binding scFv-Fc. (**E**) Assessment of agonist activity of 6-6 IgG on cells expressing LRP6 mutants. HEK293 cells were transfected with a plasmid encoding the wild-type LRP6 (WT), or the K662A/K684A double mutant, along with the Wnt3a-expression plasmid and the STF reporter expression plasmid, and incubated with or without 6-6 IgG. Error bars represent SD for n = 2. * P < 0.05. (**F**) Differentiation of critical binding sites of 6-6 from where Wnt ligand binds. LRP6-P3E3 domain (PDB: 3S8Z) was highlighted to show the binding sties of Wnt3a (*yellow;* E663, E708, H834, Y875, and M877) and 6-6 (*red;* K662 and K684).

### A novel mechanism of action: the 6-6 agonist antibody does not operate as a ligand surrogate

To map the binding site further, we first modeled the binding of 6-6 to LRP6 using a homology modeling-predicted structure for the 6-6 Fv and two known crystal structures of LRP6 (3S8Z and 4A0P), and identified several potential 6-6 contact sites (Supplemental **Figure S2A** and **S2B**). We then performed alanine scan mutagenesis at those sites and tested binding of 6-6 IgG to the LRP6 mutants by flow cytometry. We found that K662A and K684A single mutations caused a significant loss of 6-6 binding and the double mutant (K662A/K684A) caused a near complete loss of 6-6 binding (**Figure 2D**), thus identifying K662 and K684 as the critical contact sites. Consistent with the binding results, we found that 6-6-induced Wnt signaling activity was significantly decreased in HEK293 cells transfected with the double mutant (K662A/K684A) compared to the wild-type (WT) control (**Figure 2E**). These two sites are spatially distinct from Wnt3a-binding sites (**Figure 2F**), suggesting that 6-6 does not compete with ligand binding to LRP6.

Interestingly, based on the published structure of the complex of Wnt signaling inhibitor DKK1 and LRP6 (Cheng et al., 2011), we found that the 6-6 binding site does not overlap with that of the DKK1 binding site (**Figure 3A**). We thus studied binding of 6-6 to LRP6 in the presence of Wnt ligand (rhWnt3a) and inhibitor (DKK1) using biolayer interferometry. LRP6 was loaded on the biosensor, followed sequentially by binding with rhWnt3a and then 6-6 Fab or DKK1. As expected by the epitope mapping result, the 6-6 Fab bound to the LRP6-Wnt3a complex, whereas the DKK1 could not (**Figure 3B**). We next performed the STF reporter assay to evaluate if DKK1 blocks the 6-6 agonist activity. As a control, we first performed the reporter assay using rhWnt3a (**Figure 3C**) and found that Wnt signaling was inhibited by DKK1. We next performed the assay with 6-6 and found that the agonist activity persisted in the presence of DKK1 (**Figure 3D**). In addition, we sought to determine how DKK1 affects Wnt signaling when both Wnt3a and 6-6 are present. We performed the STF reporter assay using HEK293 cells transfected with the Wnt3a-expression plasmid, with or without 6-6. As shown in **Figure 3E**, DKK1 inhibited Wnt3a signaling when no 6-6 was added. When 6-6 was added, the total signal increased. The addition of DKK1 reduced the total signal but more than half remained, suggesting that DKK1 reduced Wnt3a-but not 6-6-mediated signaling.

**Figure 3.**
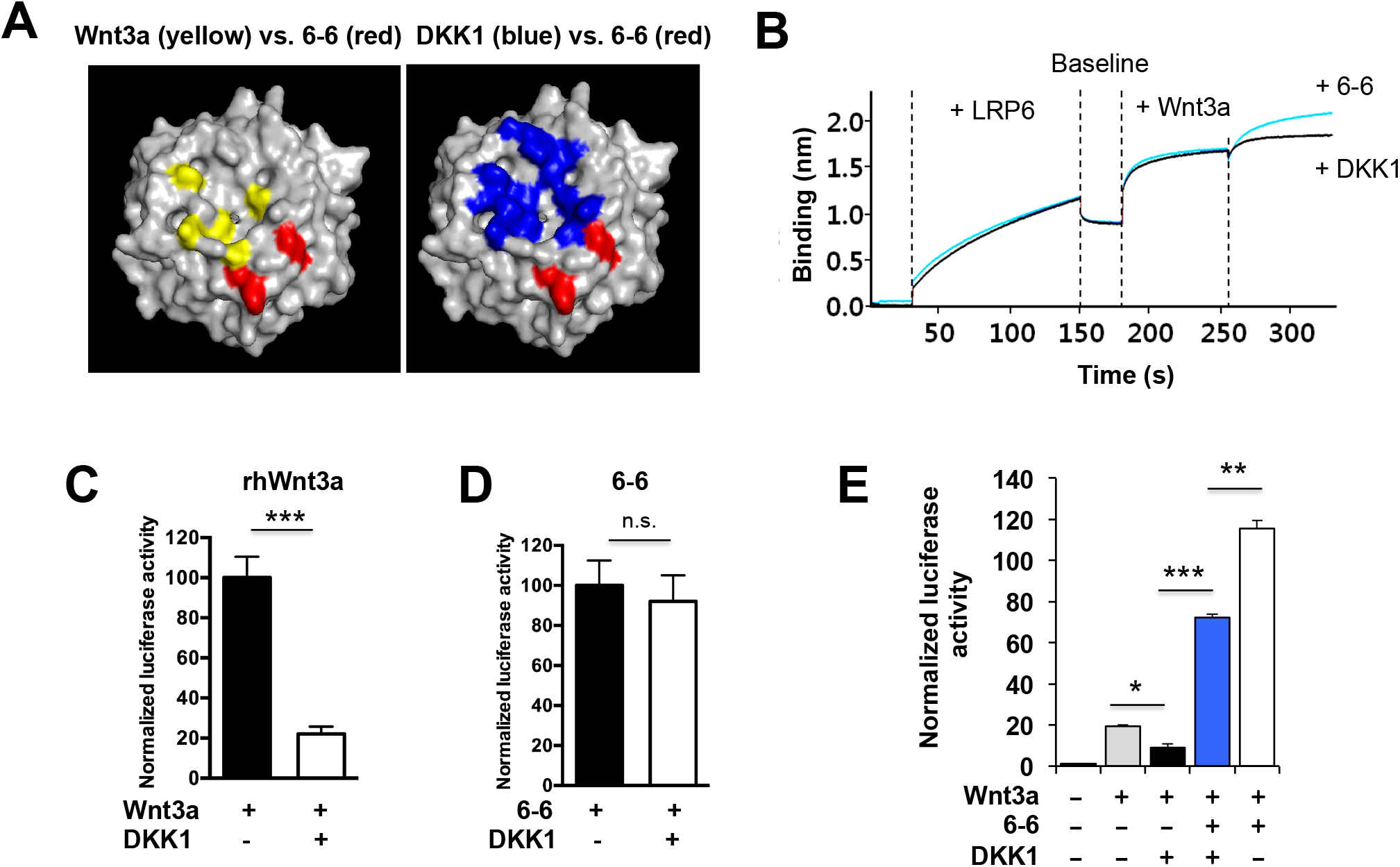
6-6 agonist effect persists in the presence of the Wnt pathway inhibitor DKK1. (**A**) Binding site comparison between DKK1 (right panel, *blue;* I681, Y706, E708, W767, R792, D811, H834, F836, W850, Y875, and M877)) and 6-6 (*red;* K662 and K684) on LRP6. The LRP6 P3 domain (631-890) is shown. To provide a reference view, the binding sites for Wnt3a (yellow) vs. 6-6 (red) is shown again in the left panel. The binding sites of Wnt3a and DKK1 overlap with each other (spatial as well as specific residues, e.g., E708, H834, Y875, and M877) but not with 6-6. (**B**) Binding kinetics of 6-6 Fab to LRP6 pre-bound by Wnt3a. The LRP6-loaded biosensor was first incubated with the rhWnt3a then a mixture of rhWnt3a and 6-6 Fab (blue line, which shows further bindin*g*), or rhWnt3a and DKK1 (as a control, black line, which shows no further binding). (**C**) Inhibition of Wnt3a signaling by DKK1. The STF reporter assay was performed on HEK293 cells with 10 nM rhWnt3a with or without 1 nM DKK1. Error bars represent SD for n = 3. *** P < 0.001. (**D**) 6-6 overcomes DKK1 inhibition. The STF reporter assay was performed on HEK293 cells with 10 nM 6-6 IgG with or without 1 nM DKK1. Error bars represent SD for n = 3. n.s.: not significant. (**E**) Effects of DKK1 on the agonist activity of 6-6 on HEK293 cells transfected with the STF reporter and the Wnt3a-expression plasmid, with or without 6-6 IgG (50 nM) or DKK1 (20 nM), as indicated. Error bars represent SD for n = 2. * P < 0.05, ** P < 0.01, *** P < 0.001.

Taken together, these results reveal a new mechanism of action of the agonist antibody 6-6: it does not operate as a ligand surrogate. Instead, it binds to a unique site on LRP6 that does not overlap with where ligands and inhibitors bind, works additively with Wnt ligands, and retains the agonist function in the presence of endogenous Wnt inhibitors.

### Canonical Wnt pathway activation by the 6-6 antibody is amplified by R-spondin

We next sought to determine if the agonist activity of 6-6 can be amplified by the Wnt signaling enhancer R-spondins (RSPOs). We performed the STF reporter assay using HEK293 cells by titrating 6-6 in the presence of RSPO2 (3 nM) without any exogenously added Wnt ligands (**Figure 4A**). For comparison, 6-6 at the highest concentration point (40 nM) without RSPO2 is used as a reference point. As shown in **Figure 4A**, the agonist effect of 6-6 IgG (no exogenous Wnt ligands added) was dramatically elevated by the addition of RSPO2. As a control, we also performed the same assay with the Wnt ligand using L cell Wnt3a-conditioned medium (Wnt3aCM) and observed a similar level of enhancement by RSPO2 (**Figure 4B**).

**Figure 4.**
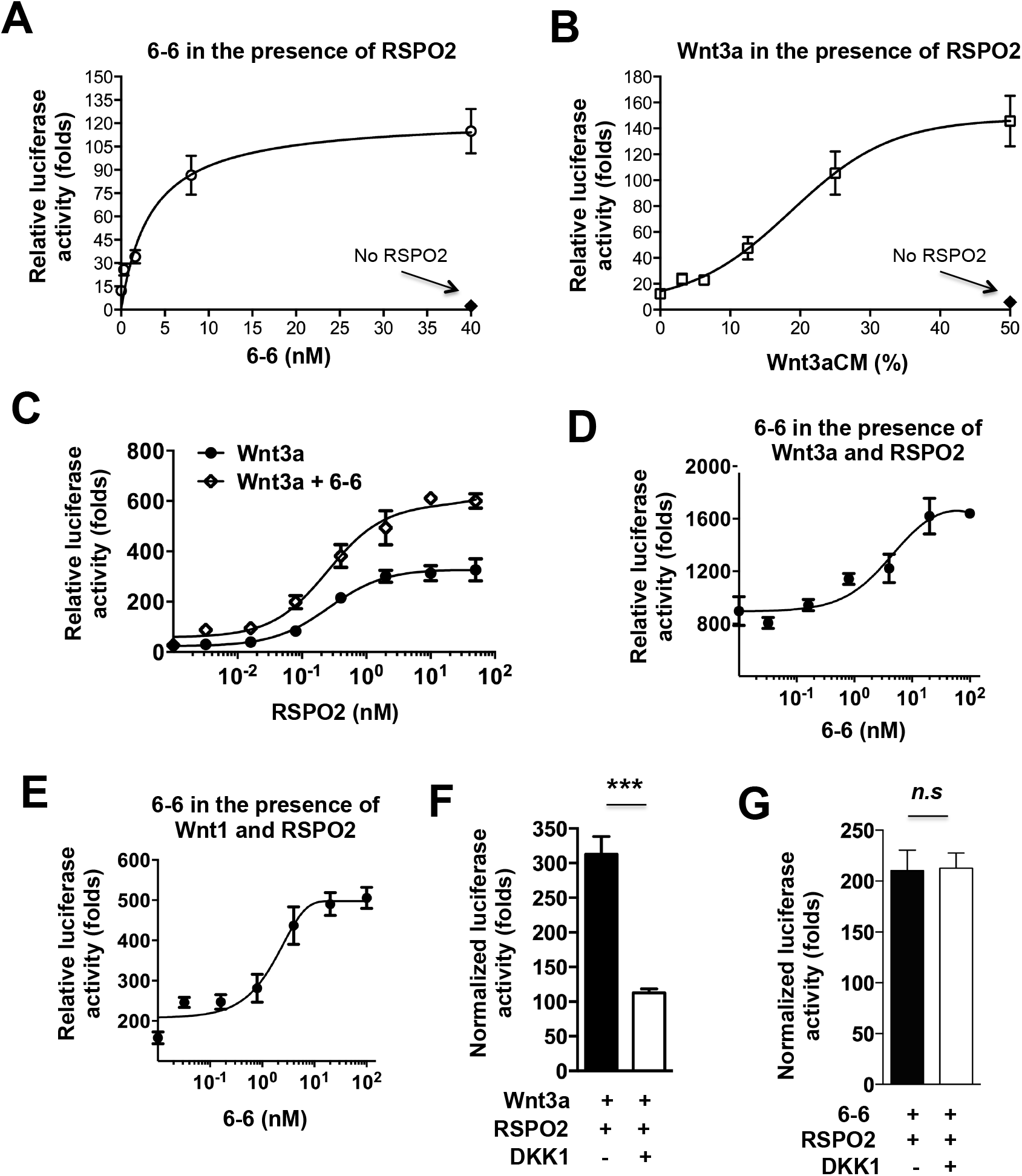
The 6-6 agonist effect is enhanced by R-spondin and persists in the presence of DKK1. (**A**) The 6-6 agonist effect is enhanced by RSPO2. The STF reporter assay was performed on HEK293 cells in the presence of 3 nM RSPO2 over a range of 6-6 IgG concentrations. For comparison, STF activity at 40 nM without RSPO2 is also shown on the graph (black diamond). The fold of enhancement with RSPO2 vs. no RSPO2 at 40 nM 6-6 IgG is 114.8 ± 14.5 over 3.98 ± 0.57 or about 29. (**B**) Wnt3a activity is enhanced by RSPO2. The STF reporter assay was performed on HEK293 cells in the presence of 3 nM RSPO2 over a range of dilutions of L cell Wnt3a conditioned media (Wnt3aCM). For comparison, STF activity at 50% Wnt3aCM without RSPO2 is also shown on the graph (black diamond). The fold of enhancement with RSPO2 vs. no RSPO2 at 50% Wnt3aCM is 145.8 ± 19.6 over 5.80 ± 0.16 or about 25. (**C**) The additive effect of 6-6 on Wnt3a signaling over a range of RSPO2 concentrations. The Wnt3a + 6-6 group: RSPO2 was titrated on HEK293 cells transfected with the STF reporter and the Wnt3a-expression construct, in the presence of a constant concentration of 6-6 IgG (20 nM). The Wnt3a group: A control RSPO2 titration without 6-6 was performed in parallel. Error bars represent SD for n = 2. (**D**) The additive agonist effect of 6-6 IgG on Wnt3a signaling over a range of 6-6 concentrations. 6-6 IgG was titrated on HEK293 cells transfected with the STF reporter and Wnt3a expression plasmids in the presence of RSPO2 (3 nM). Error bars represent SD for n = 2. (**E**) The additive agonist effect of 6-6 IgG on Wnt1 signaling over a range of 6-6 concentrations. 6-6 IgG was titrated on HEK293 cells transfected with the STF reporter and Wnt1 expression plasmids in the presence of RSPO2 (3 nM). Error bars represent SD for n = 2. (**F**) DKK1 inhibits Wnt3a/RSPO2-induced β-catenin signaling. HEK293 cells were transfected with the STF reporter and the Wnt3a-expression plasmids and incubated with RSPO2 (5 nM) or RSPO2 (5 nM) plus DKK1 (15 nM). Error bars represent SD for n = 2. *** P < 0.001. (**G**) DKK1 has no significant effects on 6-6/RSPO2-induced Wnt/β-catenin signaling enhancement. HEK293 cells transfected with the STF reporter plasmid were incubated with 6-6 IgG (100 nM), RSPO2 (5 nM), or DKK1 (15 nM). No Wnt ligands were added. Error bars represent SD for n = 2. n.s: not significant.

We next sought to determine if the additive effect of 6-6 and Wnt3a on canonical Wnt signaling (shown previously in Figure 1E where no R-spondin was present) persists in the presence of varying levels of R-spondin. We performed the STF reporter assay using Wnt3a-transfected HEK293 cells with or without 20 nM 6-6, in the presence of a range of RSPO2 concentrations. As shown in **Figure 4C**, the additive effect was observed throughout the RSPO2 concentration range. We further studied the additive effect as a function of 6-6 concentrations for two Wnt ligands, Wnt3a (**Figure 4D**) and Wnt1 (**Figure 4E**). For Wnt3a, we performed the STF reporter assay using HEK293 cells transfected with the Wnt3a expression plasmid in the presence of RSPO2 (5 nM) across a range of 6-6 concentrations, and found that 6-6 showed a concentration dependent additive effect with Wnt3a (**Figure 4D**). Similar results were obtained for Wnt1, where the STF reporter assay was performed on HEK293 cells transfected with the Wnt1 expression plasmid in the presence of RSPO2 (5 nM) across a range of 6-6 concentrations (**Figure 4E**).

Next, we studied if the R-spondin enhancement of the 6-6 agonist effect is blocked by the inhibitor DKK1. As a control, we first performed the STF reporter assay using HEK293 cells transfected with the Wnt3a-expression plasmid and incubated with RSPO2 (5 nM) or RSPO2 (5 nM) plus DKK1 (15 nM). As shown in **Figure 4F**, the R-spondin enhanced Wnt signaling was significantly reduced by DKK1. We then performed the STF reporter assay using 6-6 (100 nM) and RSPO2 (5 nM), with or without DKK1 (15 nM). As shown in **Figure 4G**, RSPO2 enhancement of 6-6 agonist effect was not significantly inhibited by DKK1.

Taken together, these data further support that 6-6 acts additively but not competitively with Wnt ligands and responds to RSPO2-mediated signaling enhancement with or without exogenously added Wnt ligands. Unlike Wnt ligands and ligand surrogates, the 6-6 agonist activity is not inhibited by endogenous inhibitors that bind to ligand binding sites.

### Biological application of Wnt agonist antibody: effects on osteoblast differentiation *in vitro*

One of the biological consequences of canonical Wnt signaling activation is the induction of osteoblast differentiation and bone formation. We investigated Wnt-agonist effect of 6-6 IgG on mouse pre-osteoblast MC3T3-E1 and bone marrow-derived mesenchymal C3H/10T1/2 cell lines. We first analyzed the cross-species binding of 6-6 to human and mouse LRP6. The 6-6 IgG bound to both human and mouse LRP6 ectodomains (**Figure 5A**). We then assessed the effect of 6-6 on Wnt/β-catenin signaling activation by STF reporter assays. Wnt3a induced reporter activity, and addition of 6-6 IgG significantly enhanced signaling in both MC3T3-E1 (**Figure 5B**) and CH3/10T1/2 (**Figure 5C**) cell lines.

**Figure 5.**
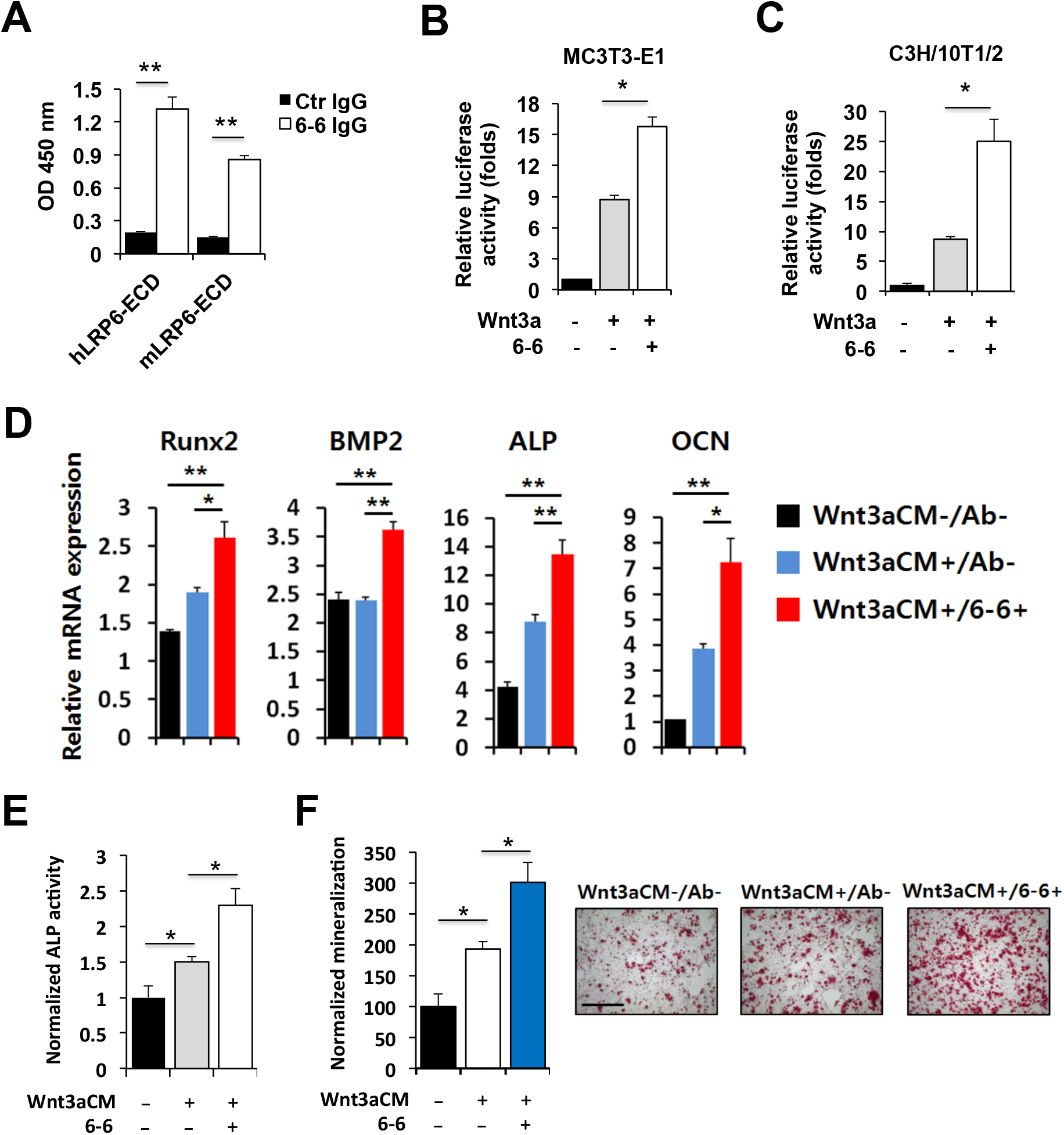
Wnt-agonist antibody promotes osteoblastic differentiation *in vitro*. (**A**) Cross-species binding of 6-6 IgG to human and mouse LRP6. Recombinant extracellular domain (ECD) of human (hLPR6-ECD) or mouse LRP6 (mLRP6-ECD) was used in an ELISA assay to assess 6-6 IgG binding. Ctrl IgG: a non-binding human IgG. Error bars represent SD for n = 2. ** P < 0.01. (**B**) and (**C**) Wnt/β-catenin signaling enhancement by 6-6 IgG in mouse cell lines. MC3T3-E1 (B) or C3H/10T1/2 (C) cell line was transfected with the Wnt3a-expression and the STF reporter plasmids and further incubated with or without 6-6 IgG. Luciferase activity was normalized against a control group transfected with the reporter construct only. Error bars represent SD for n=2. * P < 0.05. (**D**) Osteoblastic gene expression analysis. C3H/10T1/2 cells were incubated for 3 days with Wnt3aCM or 6-6 IgG as indicated. Expression of osteoblast marker genes (RUNX2, BMP2, ALP, and OCN) was assessed by qRT-PCR. Relative mRNA expression levels were calculated using the comparative Ct method and normalized to GAPDH gene. Wnt3aCM-/Ab-: no Wnt3aCM and no 6-6. Wnt3aCM+/Ab-: with Wnt3aCM but no 6-6. Wnt3aCM+/6-6+: with Wnt3a and 6-6. * P < 0.05, ** P < 0.01. (**E**) ALP activity induced by the Wnt-agonist 6-6 antibody. C3H/10T1/2 cells were cultured in Wnt3aCM with or without 6-6 IgG for 7 days. Cell lysates were used to measure ALP activity, which was normalized against a control group without Wnt3aCM and 6-6 treatment. Error bars represent SD (n = 2). * P < 0.05. (**F**) Assessment of matrix mineralization activity. C3H/10T1/2 cells were cultured for 21 days in osteogenic medium supplemented with Wnt3aCM or 6-6 IgG as indicated. Alizarin Red staining assay was applied to quantify mineralization (*left*), which was normalized against the control (Wnt3aCM-/Ab-). The measurement was presented as mean ± SD (n = 2). * P < 0.05. Representative images were shown (*right*). Scale bar: 200 μm.

We next studied induction of osteoblastic differentiation of C3H/10T1/2 cells by measuring expression of osteoblast marker genes (RUNX2, BMP2, ALP, and OCN) by qRT-PCR. As shown in **Figure 5D**, the Wnt-agonist 6-6 induced expression of Wnt downstream genes involved in osteoblastic differentiation, with 6-6 combined with Wnt3a conditioned media (Wnt3aCM) being more potent than Wnt3aCM alone. In addition to mRNA expression, we also measured alkaline phosphatase activity (ALP) in C3H/10T1/2 cells incubated with 6-6. As shown in **Figure 5E**, Wnt3aCM upregulated ALP activity, and the addition of 6-6 further increased ALP activity in the presence of Wnt3aCM.

In order to directly assess the osteoblastic commitment of C3H/10T1/2 cells, we conducted *in vitro* mineralization assay (Gregory et al., 2004) in the presence of 6-6 and Wnt3aCM. As shown in **Figure 5F**, Wnt3aCM increased mineralization that was further enhanced by the addition of 6-6. These data suggest that the Wnt-agonist antibody 6-6 promotes osteoblast differentiation and acts additively with natural Wnt ligands.

### The Wnt-agonist antibody 6-6 counteracts bone inhibitory effects by multiple myeloma cells *in vitro* and *in vivo*

Certain types of primary and metastatic cancers locate to the bone and cause extensive bone remodeling in patients. Multiple myeloma resides in the bone marrow and is known to induce extensive bone loss by secretion of inhibitors of canonical Wnt signaling (Edwards, 2008; Glass et al., 2003). Since our novel Wnt-agonist antibody 6-6 does not compete with known Wnt inhibitors for LRP6 binding, we reasoned that 6-6 could effectively counteract Wnt inhibitors produced by myeloma cells. To test this hypothesis, we studied the multiple myeloma cell line MM1.S for the presence of Wnt inhibitory activities in conditioned media on HEK293 cells co-transfected with the STF reporter and the Wnt3a-expression construct. As shown in **Figure 6A**, conditioned media from MM1.S (MM1.S-CM) showed significant inhibitory effect compared to the control (either conditioned media from the control cell line HEK293 or no conditioned media). Furthermore, in the presence of both Wnt3a and MM1.S-CM, the 6-6 IgG restored Wnt signaling inhibited by MM1.S-CM and even further stimulated signaling at higher concentrations of 6-6 (**Figure 6B**).

**Figure 6.**
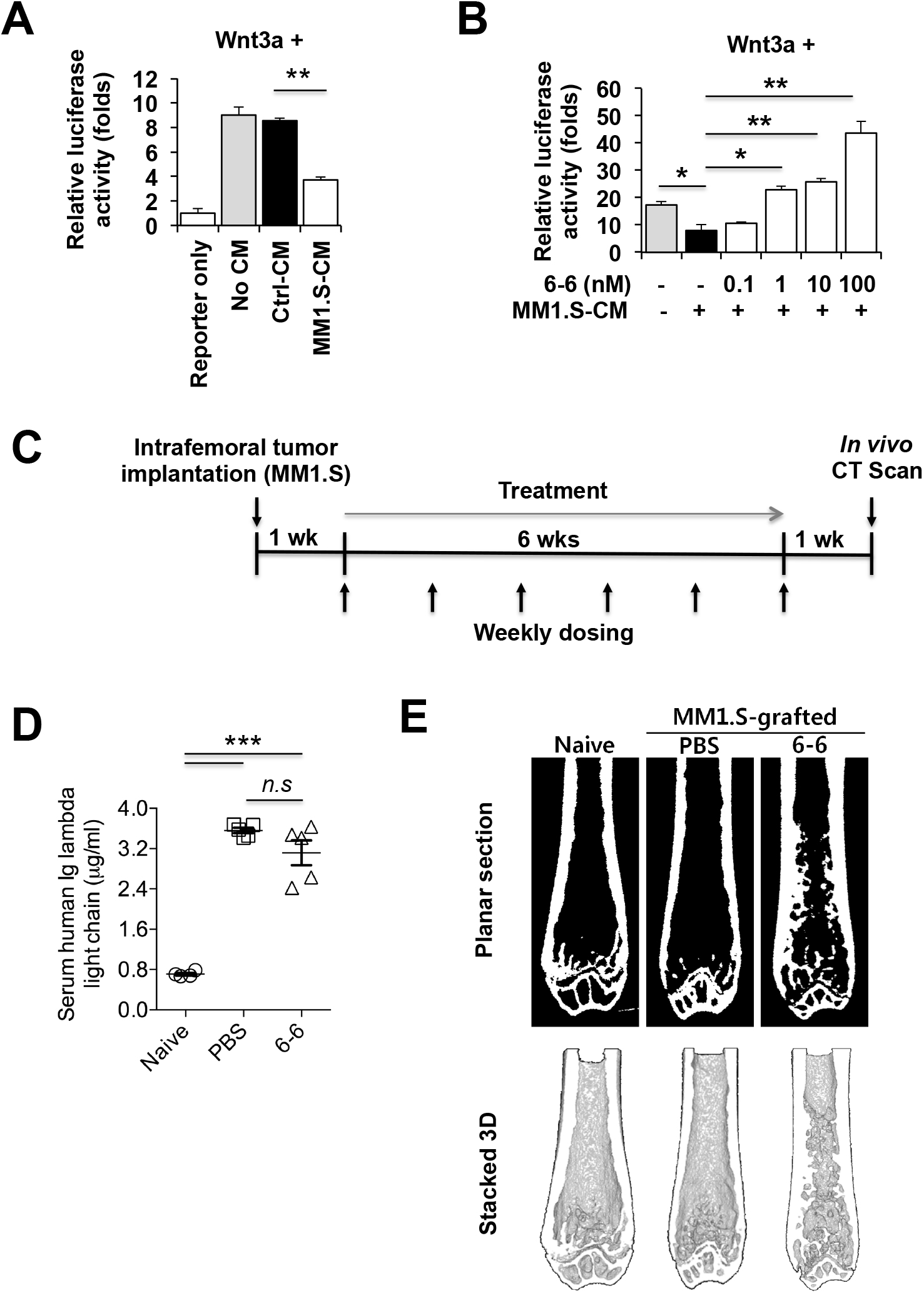
6-6 overcomes multiple myeloma-mediated Wnt signaling inhibition. (**A**) Inhibition of Wnt3a/β-catenin signaling by multiple myeloma cells. HEK293 cells transfected with the STF reporter and the Wnt3a-expression plasmids were incubated in conditioned media (CM) obtained from the multiple myeloma cell MM1.S and HEK293 (as a control, Ctrl-CM). Values represent mean ± SD for n = 2. ** P < 0.01. (**B**) 6-6 overcomes Wnt signaling inhibition caused by MM1.S-CM. HEK293 cells were transfected with the STF reporter and Wnt3a-expression constructs and incubated in MM1.S-CM with varying concentrations of 6-6 IgG. Values represent mean ± SD (n = 2). * P < 0.05, ** P < 0.01. (**C**) Outline of animal study. MM1.S cells were intrafemorally injected in the right femur and were allowed to establish for 1 week. A total of 6 weekly intraperitoneal injections of 6-6 IgG (10 mg/kg) or PBS (vehicle control) were given (n = 5/group). Femurs from live mice were scanned by micro-CT 1 week after termination of dosing. A week after *in vivo* scan, mice were sacrificed and serum and femur tissues were collected for further analysis. (**D**) Evaluation of human Ig-lambda light chain concentration in serum by ELISA. Naïve: mice without MM1.S implantation. PBS: mice with MM1.S implantation injected with PBS (vehicle control). 6-6: mice with MM1.S implantation injected with 6-6 IgG. *** P < 0.001. n.s.: not significant. (**E**) Planar and 3D views of whole femur obtained from micro-CT. Micro-CT images were used to generate clear planar sections (*top*) and to reconstruct the stacked 3D views (*bottom*).

To study the effect of 6-6 on bone remodeling *in vivo*, an intrafemoral osteolytic model was established by implantation of MM1.S cells into the right femur of NSG mice. As outlined in **Figure 6C**, antibody treatment started 1 week post implantation and continued for 6 weeks by weekly intraperitoneal (i.p.) injection. Live mice were scanned by micro-CT to assess changes of femoral bone structure. To confirm tumor establishment, we first analyzed human Ig-lambda light chain levels in serum by ELISA. As shown in **Figure 6D**, the concentration of human Ig-lambda light chain in the MM1.S-implanted groups (PBS and 6-6) was significantly higher than that of the group with no MM1.S implantation (Naive). The light chain levels in the 6-6 group were lower than those in the PBS group, but the difference did not reach statistical significance (**Figure 6D**). By immunohistochemistry study, MM1.S myeloma cells established in the right femur were also detected by anti-human Ig-lambda antibody (Supplemental **Figure S3**). The bone-forming effect of the 6-6 IgG was assessed by micro-CT analysis. Whole femurs were analyzed to generate planar and 3D images from CT-scanned data. As shown in **Figure 6E**, intrafemoral MM1.S implantation resulted in bone loss especially in the trabecular bone area (Naive vs. PBS). Strikingly, 6-6 treatment reversed the femoral bone loss compared to the control (6-6 IgG vs. PBS, **Figure 6E**), indicating that the Wnt-agonist antibody 6-6 promotes bone formation *in vivo*.

We further investigated quantitatively bone formation induced by 6-6 IgG. Region of interest (ROI) to assess bone structures was designated as shown in **Figure 7A**, and micro-CT images were reconstructed for 3D view and quantification of bone micro-architectures. By analyzing trabecular ROI, compared to the Naive group with no myeloma cell implantation, trabecular bone in the vehicle control group (PBS) was catabolized by implanted MM1.S cells (**Figure 7B**). However, consistent with the whole femur image analysis, 6-6 IgG treatment showed trabecular-anabolic activity (**Figure 7B**). 6-6 IgG treatment resulted in a significant increase in bone volume over tissue volume (BV/TV) and trabecular bone thickness (Tb.Th) (**Figure 7C** and **7D**, respectively). In addition to the distal femur, we analyzed cortical bone in the proximal femur legion. By reconstructing the cortical ROI and measuring cortical bone thickness (Ct.Th), we found that the 6-6 IgG significantly increased cortical bone formation compared to the control group (PBS) (**Figure 7E** and **7F**), suggesting that the 6-6 Wnt-agonist effect stimulates both trabecular and cortical bone formation. Taken together, the novel Wnt-agonist antibody 6-6 reverses bone loss in the intrafemoral MM1.S myeloma model, demonstrating its potential in treating osteolytic diseases.

**Figure 7.**
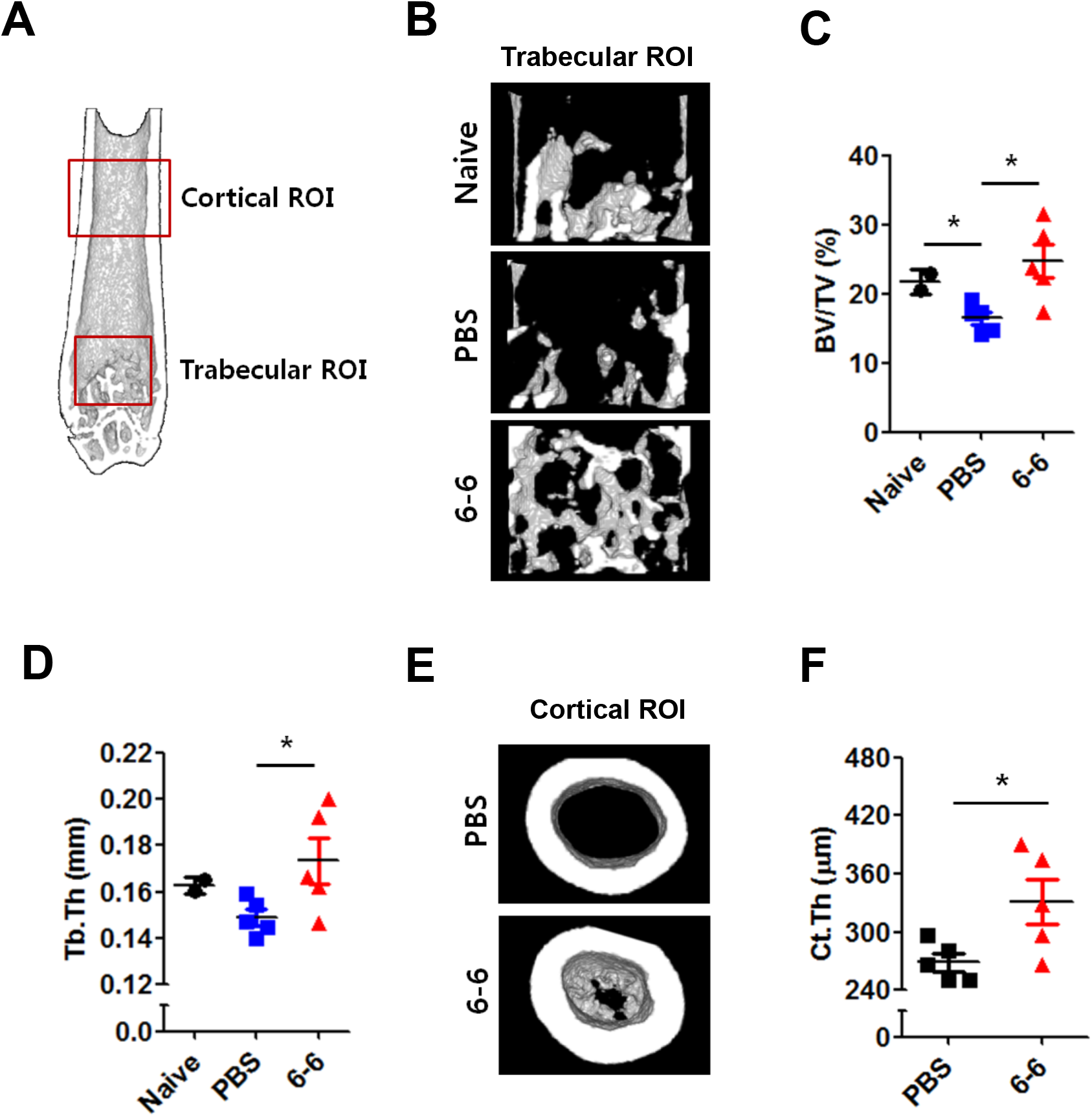
Wnt agonist antibody reverses bone loss in the intrafemoral MM1.S model. (**A**) Designated region of interest (ROI) in femurs. Micro-CT images are used to reconstruct 3D data set of trabecular and cortical bone regions as indicated. (**B**) 6-6-induced restoration of trabecular bone loss. The micro-CT images of the naive or MM1.S-implanted femurs (PBS and 6-6 IgG) were used to reconstruct 3D architectures of trabecular bone regions. Representative images were shown. (**C**) and (**D**) Quantification of trabecular bone microarchitectures. Bone volume over tissue volume (BV/TV, panel C) and trabecular bone thickness (Tb.Th, panel D) were quantified to compare bone formation activity between the groups. * P < 0.05. (**E**) Enhancement of cortical bone formation by 6-6. Micro-CT images of cortical bone were reconstructed from proximal femur regions. Representative images were shown. (**F**) Cortical bone thickness (Ct.Th) was measured and compared (PBS vs. 6-6). * P < 0.05.

## Discussion

Due to the importance of the Wnt signaling pathway in regenerative medicine, there has been a growing interest to develop therapeutics that activate this pathway. Wnt ligands, however, are post-translationally modified (e.g., lipidation) and difficult to produce as recombinant biologic drugs. Several studies have focused on the ligand surrogate approach, where the design principle is to mimic natural ligand-receptor interaction with part of the ligand complex being replaced by a receptor-binding antibody fragment or recombinant protein, which can be further modified to create multivalent interactions to increase potency by crosslinking the Wnt receptor complex (Chen et al., 2020; Fowler et al., 2021; Gong et al., 2010; Janda et al., 2017; Luca et al., 2020; Tao et al., 2019). A challenge for the ligand surrogate approach is that the ligand surrogate binds to the same area where endogenous activators and inhibitors bind, making it challenging to not compete with Wnt ligands and/or retain the agonist activity in the presence of inhibitors. Our study addresses those challenges and identifies a new type of agonists that does not follow the ligand surrogate design. **Figure 8** summarizes key differences between these two mechanisms of action.

**Figure 8.**
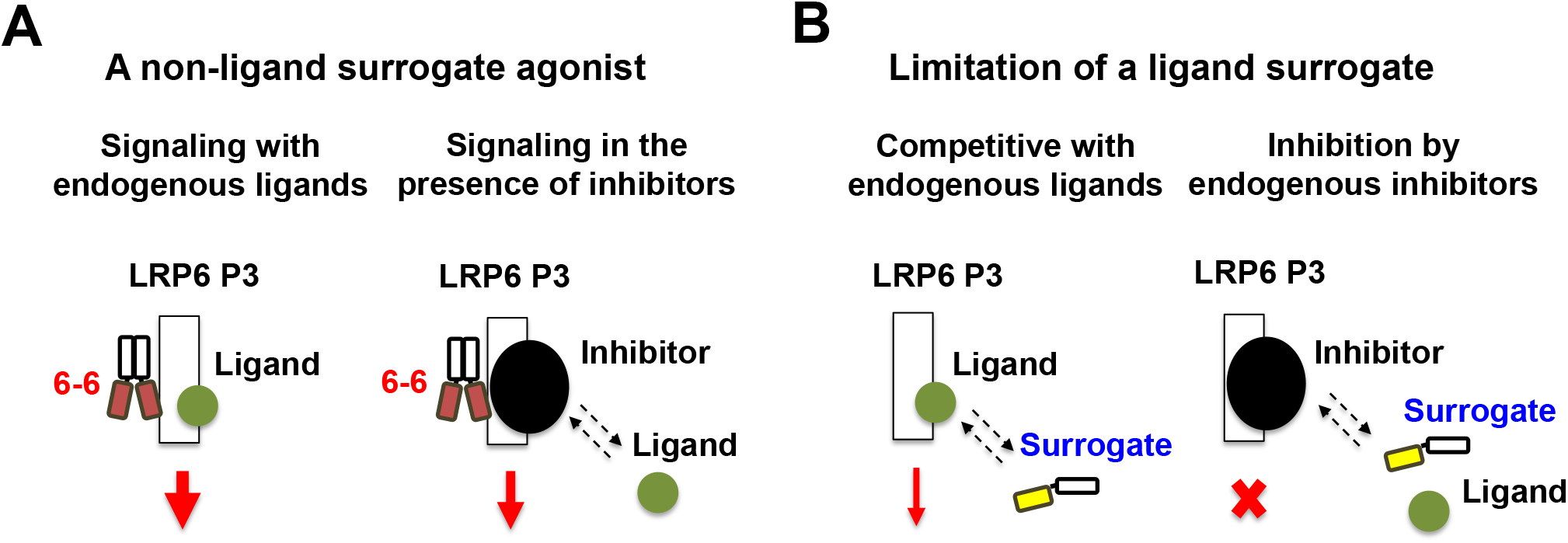
Summary of key differences between our non-ligand surrogate agonist vs. a ligand surrogate. As shown in (**A**), 6-6 does not bind to where known endogenous ligands and inhibitors bind, works additively with endogenous ligands, and is able to activate Wnt signaling in the presence of inhibitors. In contrast, as shown in (**B**), a ligand surrogate is subject to competitive binding by both endogenous ligands and inhibitors, and is ineffective in activating the Wnt pathway when inhibitors are present. Dashed lines indicate competition by inhibitors. For simplicity, only the LRP6 P3 domain is drawn; other components of the Wnt receptor complex are not drawn.

We discovered a novel Wnt-agonist human monoclonal antibody 6-6 that is capable of activating Wnt signaling even when Wnt ligands are not provided. The addition of Wnt ligands further enhanced the agonist effect of 6-6. This agonist antibody binds the P3 domain of LRP6. Our epitope mapping study further defined critical binding sites for 6-6 that do not overlap with Wnt ligand binding sites. In addition, 6-6 responds to R-spondin, achieving enhanced Wnt signaling activation similar to that of Wnt ligand in the presence of R-spondin. Importantly, unlike a ligand surrogate, the 6-6 Wnt agonist effect is not significantly inhibited by endogenous Wnt inhibitors such as DKK1. This marks a major distinction from previous Wnt activator studies that have been focused on developing ligand surrogates, either in monovalent or multivalent forms (Chen et al., 2020; Fowler et al., 2021; Janda et al., 2017; Tao et al., 2019).

A known biological effect of canonical Wnt signaling is bone formation. Therefore we sought to develop Wnt agonist-based therapeutics in the area of pathological bone loss induced either by cancer or aging. In this study we focused on multiple myeloma-induced bone loss. Multiple myeloma cells reside in the bone marrow and inhibit Wnt signaling in osteoblast cells either directly by secreting the Wnt inhibitor DKK1 or indirectly by stimulating the production of Wnt inhibitor SOST in the bone microenvironment (Delgado-Calle et al., 2016; Qiang et al., 2008). We found that our Wnt agonist antibody treatment promotes osteoblastic differentiation *in vitro* and reverses myeloma-induced bone loss *in vivo*, suggesting a therapeutic application of this antibody in restoring bone loss in multiple myeloma patients following anti-cancer treatment. Further studies will determine if this novel agonist antibody also has therapeutic potential in restoring bone loss induced by other cancer types, by aging in postmenopausal osteoporosis patients, and in rare genetic diseases such as osteogenesis imperfecta.

The unique mechanism of action of our Wnt agonist antibody also differentiates it from current anti-osteoporosis drugs that targets the Wnt pathway such as romosozumab, an FDA approved humanized monoclonal antibody that blocks SOST and shows clinical activity of improving bone formation in osteoporosis patients (Lewiecki et al., 2018; Markham, 2019; Sølling et al., 2018). This is an anti-inhibitory ligand approach and has a few limitations. If the endogenous Wnt ligand is not present in sufficient amounts, blocking SOST will have marginal effects. In addition, inhibitors other than SOST are not subject to neutralization by romosozumab. In contrast, our Wnt agonist antibody 6-6 can stimulate canonical Wnt signaling in Wnt ligand-low or insufficient setting, and overcomes inhibition by Wnt inhibitors. Moreover, the agonist effect of 6-6 is further enhanced by R-spondin, an endogenous amplifier of Wnt/β-catenin signaling, with or without Wnt ligands added. These features of our agonist antibody could offer broad applicability for degenerative bone lesions, especially in conditions where endogenous Wnt ligands are insufficient or secretion of Wnt inhibitors is increased either in quantity or variety.

In summary, our study uncovered a canonical Wnt agonist antibody that does not operate as a ligand surrogate. It binds to a unique site on LRP6 that does not overlap with where known endogenous Wnt ligands or inhibitors bind. This novel agonist antibody showed abilities to activate canonical Wnt pathway in the presence of inhibitors and when no Wnt ligands are provided, and to restore bone loss *in vivo* in a cancer-induced osteolytic legion model. This unique mechanism of action differentiates it from both ligand surrogates and anti-inhibitors such as romosozumab, opening a novel path for therapeutic development against bone loss brought on by cancer or aging and other degenerative lesions driven by insufficient Wnt signaling.

## Acknowledgement

We thank Tony L. Huynh for help with in vivo micro-CT scanning; Byron C. Hann, Veronica Steri, Paul Phojanakong, Donghui Wang, and Fernando Salangsang at the UCSF Preclinical Therapeutic Core for help with intra-femur tumor injection and antibody dosing. The work was supported in part by the National Institutes of Health (R01 CA118919, CA129491, and CA171315 to BL), and a fellowship grant to NKL from the Basic Science Research Program of the National Research Foundation of Korea, the Ministry of Education, Science and Technology (2013R1A6A3A03060495).

## Reference

Baron, R., and Kneissel, M. (2013). WNT signaling in bone homeostasis and disease: from human mutations to treatments. Nature Medicine 19, 179–192.

Chen, H., Lu, C., Ouyang, B., Zhang, H., Huang, Z., Bhatia, D., Lee, S.J., Shah, D., Sura, A., Yeh, W.C., et al. (2020). Development of Potent, Selective Surrogate WNT Molecules and Their Application in Defining Frizzled Requirements. Cell chemical biology 27, 598–609 e594.

Chen, J.X., Yan, H.W., Ren, D.N., Yin, Y., Li, Z., He, Q.Q., Wo, D., Ho, M.S.C., Chen, Y.H., Liu, Z.M., et al. (2014). LRP6 dimerization through its LDLR domain is required for robust canonical Wnt pathway activation. Cellular Signalling 26, 1068–1074.

Chen, S., Bubeck, D., MacDonald, B.T., Liang, W.X., Mao, J.H., Malinauskas, T., Llorca, O., Aricescu, A.R., Siebold, C., He, X., et al. (2011). Structural and Functional Studies of LRP6 Ectodomain Reveal a Platform for Wnt Signaling. Developmental Cell 21, 848–861.

Cheng, Z., Biechele, T., Wei, Z., Morrone, S., Moon, R.T., Wang, L., and Xu, W. (2011). Crystal structures of the extracellular domain of LRP6 and its complex with DKK1. Nature structural & molecular biology 18, 1204–1210.

Clevers, H., Loh, K.M., and Nusse, R. (2014). An integral program for tissue renewal and regeneration: Wnt signaling and stem cell control. Science 346, 54–+.

Delgado-Calle, J., Anderson, J., Cregor, M.D., Hiasa, M., Chirgwin, J.M., Carlesso, N., Yoneda, T., Mohammad, K.S., Plotkin, L.I., Roodman, G.D., et al. (2016). Bidirectional Notch Signaling and Osteocyte-Derived Factors in the Bone Marrow Microenvironment Promote Tumor Cell Proliferation and Bone Destruction in Multiple Myeloma. Cancer Research 76, 1089–1100.

Doube, M., Klosowski, M.M., Arganda-Carreras, I., Cordelieres, F.P., Dougherty, R.P., Jackson, J.S., Schmid, B., Hutchinson, J.R., and Shefelbine, S.J. (2010). BoneJ Free and extensible bone image analysis in ImageJ. Bone 47, 1076–1079.

Edwards, C.M. (2008). Wnt signaling: bone’s defense against myeloma. Blood 112, 216–217.

Ettenberg, S.A., Charlat, O., Daley, M.P., Liu, S., Vincent, K.J., Stuart, D.D., Schuller, A.G., Yuan, J., Ospina, B., Green, J., et al. (2010). Inhibition of tumorigenesis driven by different Wnt proteins requires blockade of distinct ligand-binding regions by LRP6 antibodies. Proceedings of the National Academy of Sciences of the United States of America 107, 15473–15478.

Florio, M., Gunasekaran, K., Stolina, M., Li, X.D., Liu, L., Tipton, B., Salimi-Moosavi, H., Asuncion, F.J., Li, C.Y., Sun, B.H., et al. (2016). A bispecific antibody targeting sclerostin and DKK-1 promotes bone mass accrual and fracture repair. Nature communications 7, 11505.

Fowler, T.W., Mitchell, T.L., Janda, C.Y., Xie, L., Tu, S., Chen, H., Zhang, H., Ye, J., Ouyang, B., Yuan, T.Z., et al. (2021). Development of selective bispecific Wnt mimetics for bone loss and repair. Nature communications 12, 3247.

Fulciniti, M., Tassone, P., Hideshima, T., Vallet, S., Nanjappa, P., Ettenberg, S.A., Shen, Z., Patel, N., Tai, Y.T., Chauhan, D., et al. (2009). Anti-DKK1 mAb (BHQ880) as a potential therapeutic agent for multiple myeloma. Blood 114, 371–379.

Glass, D.A., Patel, M.S., and Karsenty, G. (2003). A new insight into the formation of osteolytic lesions in multiple myeloma. New England Journal of Medicine 349, 2479–2480.

Gong, Y., Bourhis, E., Chiu, C., Stawicki, S., DeAlmeida, V.I., Liu, B.Y., Phamluong, K., Cao, T.C., Carano, R.A., Ernst, J.A., et al. (2010). Wnt isoform-specific interactions with coreceptor specify inhibition or potentiation of signaling by LRP6 antibodies. PloS one 5, e12682.

Gregory, C.A., Gunn, W.G., Peister, A., and Prockop, D.J. (2004). An Alizarin red-based assay of mineralization by adherent cells in culture: comparison with cetylpyridinium chloride extraction. Analytical Biochemistry 329, 77–84.

Iyer, S.P., Beck, J.T., Stewart, A.K., Shah, J., Kelly, K.R., Isaacs, R., Bilic, S., Sen, S., and Munshi, N.C. (2014). A Phase IB multicentre dose-determination study of BHQ880 in combination with anti-myeloma therapy and zoledronic acid in patients with relapsed or refractory multiple myeloma and prior skeletal-related events. British journal of haematology 167, 366–375.

Janda, C.Y., Dang, L.T., You, C.J., Chang, J.L., de Lau, W., Zhong, Z.D.A., Yan, K.S., Marecic, O., Siepe, D., Li, X.N., et al. (2017). Surrogate Wnt agonists that phenocopy canonical Wnt and beta-catenin signalling. Nature 545, 234–+.

Joiner, D.M., Ke, J., Zhong, Z., Xu, H.E., and Williams, B.O. (2013). LRP5 and LRP6 in development and disease. Trends in endocrinology and metabolism: TEM 24, 31–39.

Lee, N.K., Zhang, Y., Su, Y., Bidlingmaier, S., Sherbenou, D.W., Ha, K.D., and Liu, B. (2018). Cell-type specific potent Wnt signaling blockade by bispecific antibody. Scientific reports 8, 766.

Lewiecki, E.M., Blicharski, T., Goemaere, S., Lippuner, K., Meisner, P.D., Miller, P.D., Miyauchi, A., Maddox, J., Chen, L., and Horlait, S. (2018). A Phase III Randomized Placebo-Controlled Trial to Evaluate Efficacy and Safety of Romosozumab in Men With Osteoporosis. The Journal of clinical endocrinology and metabolism 103, 3183–3193.

Lien, W.H., and Fuchs, E. (2014). Wnt some lose some: transcriptional governance of stem cells by Wnt/beta-catenin signaling. Genes & Development 28, 1517–1532.

Liu, H., Liu, Z.Q., Du, J., He, J., Lin, P., Amini, B., Starbuck, M.W., Novane, N., Shah, J.J., Davis, R.E., et al. (2016). Thymidine phosphorylase exerts complex effects on bone resorption and formation in myeloma. Science Translational Medicine 8.

Luca, V.C., Miao, Y., Li, X., Hollander, M.J., Kuo, C.J., and Garcia, K.C. (2020). Surrogate R-spondins for tissue-specific potentiation of Wnt Signaling. PloS one 15, e0226928.

Markham, A. (2019). Romosozumab: First Global Approval. Drugs 79, 471–476.

McDonald, M.M., Reagan, M.R., Youlten, S.E., Mohanty, S.T., Seckinger, A., Terry, R.L., Pettitt, J.A., Simic, M.K., Cheng, T.L., Morse, A., et al. (2017). Inhibiting the osteocyte-specific protein sclerostin increases bone mass and fracture resistance in multiple myeloma. Blood 129, 3452–3464.

Munshi, N.C., and Anderson, K.C. (2013). New strategies in the treatment of multiple myeloma. Clinical cancer research: an official journal of the American Association for Cancer Research 19, 3337–3344.

Pierce, B.G., Wiehe, K., Hwang, H., Kim, B.H., Vreven, T., and Weng, Z.P. (2014). ZDOCK server: interactive docking prediction of protein-protein complexes and symmetric multimers. Bioinformatics 30, 1771–1773.

Pozzi, S., Fulciniti, M., Yan, H., Vallet, S., Eda, H., Patel, K., Santo, L., Cirstea, D., Hideshima, T., Schirtzinge, L., et al. (2013). In vivo and in vitro effects of a novel anti-Dkk1 neutralizing antibody in multiple myeloma. Bone 53, 487–496.

Qiang, Y.W., Barlogie, B., Rudikoff, S., and Shaughnessy, J.D. (2008). Dkk1-induced inhibition of Wnt signaling in osteoblast differentiation is an underlying mechanism of bone loss in multiple myeloma. Bone 42, 669–680.

Schindelin, J., Arganda-Carreras, I., Frise, E., Kaynig, V., Longair, M., Pietzsch, T., Preibisch, S., Rueden, C., Saalfeld, S., Schmid, B., et al. (2012). Fiji: an open-source platform for biological-image analysis. Nature Methods 9, 676–682.

Smith, K., Garman, L., Wrammert, J., Zheng, N.Y., Capra, J.D., Ahmed, R., and Wilson, P.C. (2009). Rapid generation of fully human monoclonal antibodies specific to a vaccinating antigen. Nature Protocols 4, 372–384.

Sølling, A.S.K., Harsløf, T., and Langdahl, B. (2018). The clinical potential of romosozumab for the prevention of fractures in postmenopausal women with osteoporosis. Therapeutic advances in musculoskeletal disease 10, 105–115.

Steinhart, Z., and Angers, S. (2018). Wnt signaling in development and tissue homeostasis. Development 145.

Su, Y., Liu, Y., Behrens, C.R., Bidlingmaier, S., Lee, N.K., Aggarwal, R., Sherbenou, D.W., Burlingame, A.L., Hann, B.C., Simko, J.P., et al. (2018). Targeting CD46 for both adenocarcinoma and neuroendocrine prostate cancer. JCI insight 3.

Tao, Y., Mis, M., Blazer, L., Ustav, M.J., Steinhart, Z., Chidiac, R., Kubarakos, E., O’Brien, S., Wang, X., Jarvik, N., et al. (2019). Tailored tetravalent antibodies potently and specifically activate Wnt/Frizzled pathways in cells, organoids and mice. eLife 8.

Weitzner, B.D., Jeliazkov, J.R., Lyskov, S., Marze, N., Kuroda, D., Frick, R., Adolf-Bryfogle, J., Biswas, N., Dunbrack, R.L., and Gray, J.J. (2017). Modeling and docking of antibody structures with Rosetta. Nature Protocols 12, 401–416.

Zhong, Z.D.A., Ethen, N.J., and Williams, B.O. (2016). Use of Primary Calvarial Osteoblasts to Evaluate the Function of Wnt Signaling in Osteogenesis. Wnt Signaling: Methods and Protocols 1481, 119–125.

