## Supplemental materials for "A non-ligand surrogate agonist antibody that enhances canonical Wnt signaling and bone regeneration"

**Supplemental Figure S1. Analysis of 6-6 binding to LRP6.** (**A**) Apparent binding affinity of 6-6 IgG. HEK293 cells were incubated with varying concentrations of 6-6 IgG, and cell binding was analyzed by flow cytometry. Apparent $K_D$ (~ 5.0 nM) was estimated by curve fitting. MFI: Mean fluorescence intensity. (**B**) The agonist 6-6 binds to a different site on LRP6-P3E3P4E4 than an antagonist antibody (E34N19) identified previously (Lee et al., 2018) that binds to the P3E3P4E4 domain. HEK293 cells were transfected with LRP6 expression plasmid and incubated with 6-6 or E34N19 scFv-phage for 1 h, in the presence of E34N19 IgG (as a competitor). Bound phages were detected by flow cytometry. MFI: Median fluorescence intensity.

**Supplemental Figure S2. Modeling 6-6 and LRP6-P3E3 interaction.** The variable fragment consisting of VH and VL of 6-6 antibody (6-6 Fv) modeled by RosettaAntibody module and the LRP6-P3E3 domain (PDB: 3S8Z, **S2A**; or PDB: 4A0P, **S2B**) were used for structural docking by ZDOCK. Blue: CDRHs of 6-6 Fv; Light grey: LRP6-P3E3 domain; Purple: Predicted potential binding sites (within 4 angstrom) by 6-6 Fv according to the 3S8Z model (shown in **S2A**, T659, G660, K662, L683, K684, T685,

H698, V699, E701, F702, G703, D735, G736, Q737, H738, R739) or the 4A0P model (shown in **S2B**, T659, G660, V661, K662, S682, L683, K684, T685, S687, H698, V699, V700, E701, F702, D735, G736, Q737, H738, R739); Yellow: Residues involved in Wnt3a binding (E663, E708, H834, Y875, M877) (Chen et al., 2011).

**Supplemental Figure S3. Immunohistochemical analysis of femoral MM1.S tumor burden**. Femur tissues obtained from PBS- or 6-6-treated mice were formalin-fixed and paraffin embedded, sectioned at 4 μm and stained with anti-human Ig-lambda (λ) light chain antibody to mark MM1.S cells, and counter-stained by hematoxylin. Distal region of representative left (no tumor implantation) and right femurs (with MM1.S injected) was shown. Brown color indicates positive human Ig-lambda staining. Scale bar = 100 μm.

**Reference:**

Chen, S., Bubeck, D., MacDonald, B.T., Liang, W.X., Mao, J.H., Malinauskas, T., Llorca, O., Aricescu, A.R., Siebold, C., He, X., *et al.* (2011). Structural and Functional Studies of LRP6 Ectodomain Reveal a Platform for Wnt Signaling. Developmental Cell *21*, 848-861.

Lee, N.K., Zhang, Y., Su, Y., Bidlingmaier, S., Sherbenou, D.W., Ha, K.D., and Liu, B. (2018). Cell-type specific potent Wnt signaling blockade by bispecific antibody. Scientific reports *8*, 766.

### Supplemental Figure S1

A

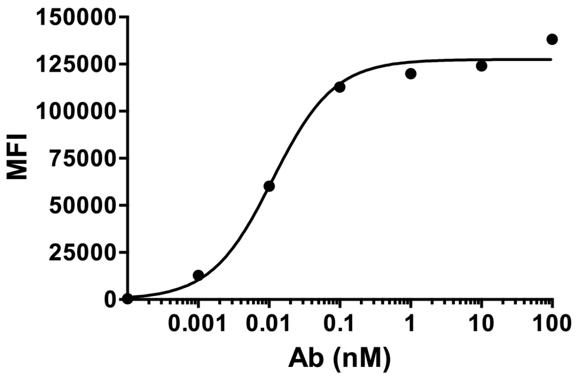


B

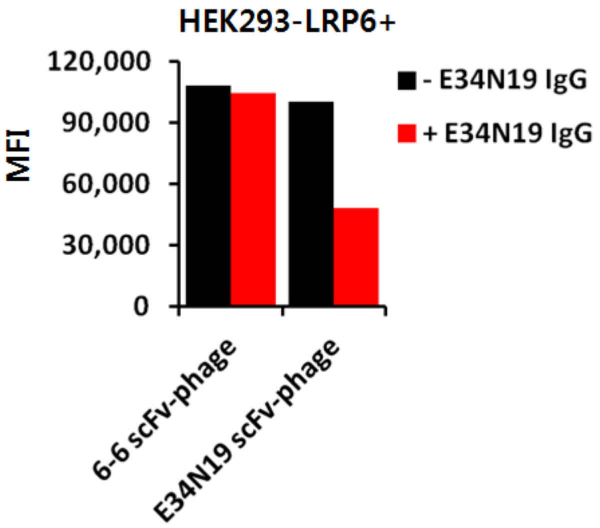

### Supplemental Figure S2

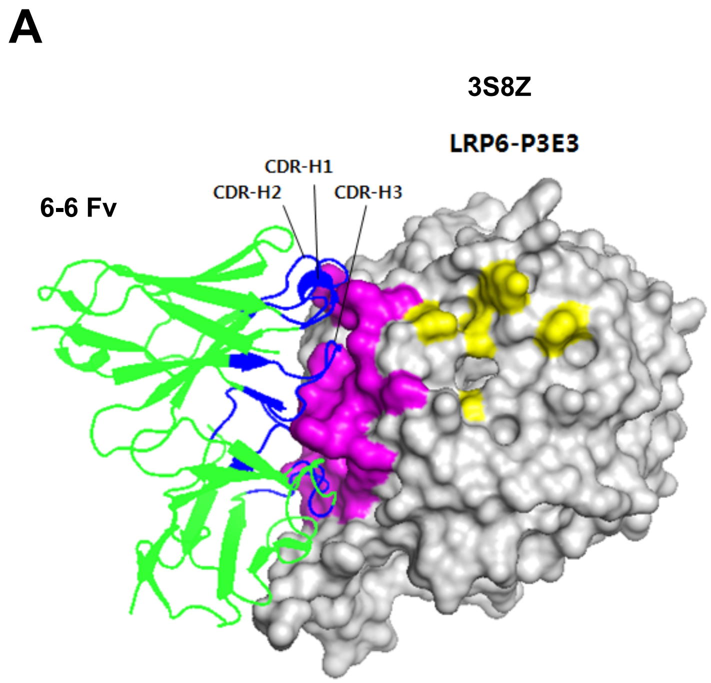


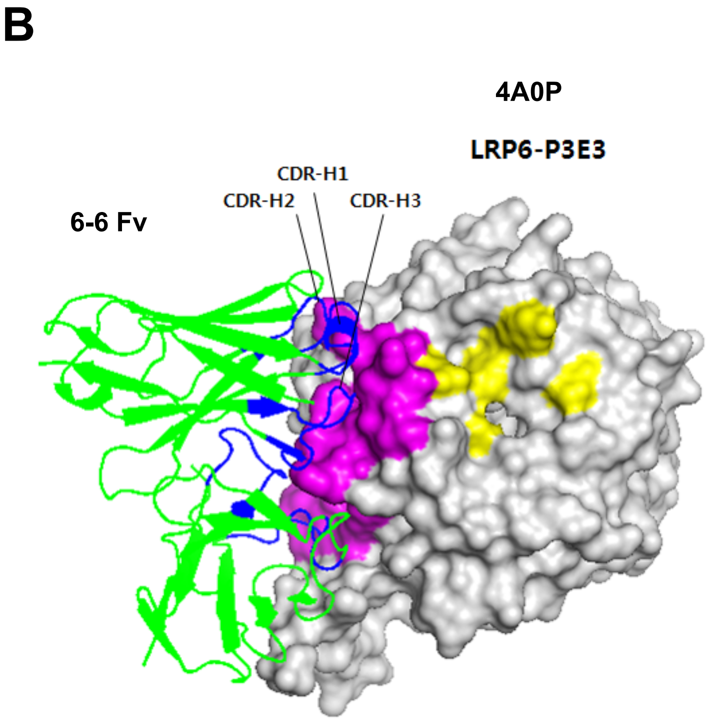

### Supplemental Figure S3

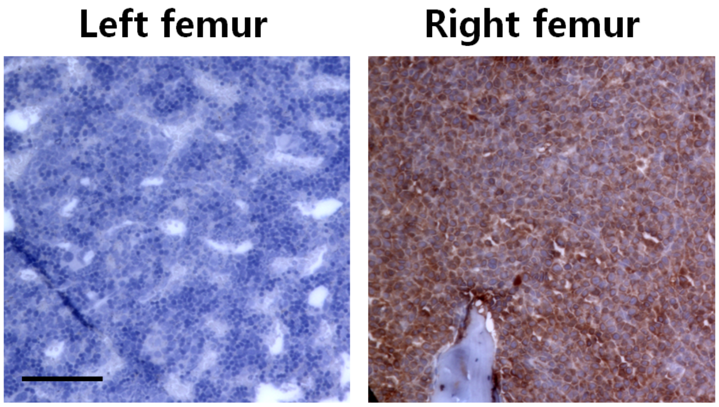
